# Multi-foci replication domains in *Haloferax volcanii* visualised by super-resolution microscopy

**DOI:** 10.64898/2026.09.02.748831

**Authors:** Dorian Noury, Titouan d’Yvoire, Ulrike Endesfelder, Stephane Duigou, Nicolas Olivier, Roxane Lestini

## Abstract

The archaeon *Haloferax volcanii* is a unique model organism to investigate replication dynamics and dissect the interplay between multiple origins, alternative replication pathways, and polyploidy. The combination of variable chromosome copy number and asynchronous replication across the cell population gives rise to pronounced phenotypic heterogeneity, further increasing system complexity. To investigate these mechanisms, we implemented (spt-)PALM and STORM microscopy to probe replication dynamics in different cell lines and growth conditions. We established the first STORM-based super-resolution imaging approach of a protein in *Haloferax volcanii*. Altogether, our results support a model in which each *H. volcanii* cell replicates a constant fraction of its chromosome copies, irrespective of total ploidy. Consequently, cells with a higher number of chromosomes exhibit proportionally more replication forks, resulting in pronounced ploidy heterogeneity across the population. Moreover, we show that replication foci are not randomly distributed but cluster into discrete Replication Domains containing multiple active sites. This organization persists across different growth conditions and is independent of canonical replication origins, as demonstrated in a strain lacking all four origins. Our findings suggest that replication coordination in *H. volcanii* is governed by spatial architecture rather than origin-dependent initiation, providing new insights into the evolutionary diversity of replication control mechanisms.

## 1 Introduction

DNA replication is a fundamental process that ensures the faithful transmission of genetic information across generations. The fundamental mechanisms of genome duplication, including origin activation, replication fork progression, and retention of replication fidelity, remain strikingly conserved across all domains of life. However, the number, spatial organisation, and regulation of replication origins are notably different between eukaryotes, bacteria, and archaea.

Most bacteria possess a single circular chromosome with a unique origin of replication. Bacterial replisomes advance at several tens of kilobases per minute (60 kb/min in *Escherichia coli* [O’Donnell et al., 2013], 40 kb/min in *Bacillus subtilis* [Pham et al., 2013]). Individual replisomes traverse long tracks — on the order of 1–2 megabases (Mb) in *E. coli* - before converging at the terminus. In contrast, eukaryotic chromosomes are vastly larger than those of prokaryotes, and their replication relies on the firing of hundreds to thousands of licensed origins. In eukaryotes, the inter-origin distance typically ranges from 10 to 200 kilobases depending on the organism and the chromosomal region. For instance, in yeast, origins are spaced approximately every 50 kilobases, while in mammalian cells, this distance can exceed 100–200 kilobases [Petryk et al., 2016, Hennion et al., 2020]. This variability reflects differences in origin licensing, chromatin accessibility, and adaptation of replication dynamics to cellular demands such as stress responses. Moreover, the organization of replication extends beyond eukaryotic origin spacing. Origins are clustered into replication domains: large chromosomal regions spanning hundreds of kilobases to megabases where origin firing is temporally coordinated. Early-replicating domains are enriched in transcriptionally active genes, while late-replicating domains often coincide with heterochromatin. This hierarchical organization ensures the complete and orderly duplication of the genome within the constrained timeframe of S phase, as observed by super-resolution microscopy [Baddeley et al., 2010, Su et al., 2020].

Archaea occupy an intermediate position: Their circular chromosomes are comparable in size to those of bacteria but may harbor either one single or multiple replication origins, depending on the species. This distribution of multiple origins appears scattered across the archaeal phylogenetic tree and shows no clear correlation with chromosome size. Comparative genomics suggests that additional origins arose independently in several lineages rather than through gradual vertical inheritance [Ausiannikava et al., 2018, Robinson and Bell, 2007, Wu et al., 2012]. Members of the TACK superphylum frequently possess chromosomes with multiple active origins, with the exception of members of the *Thaumarchaeota* phylum which possess chromosomes with a single replication origin. Pioneering work revealed that *Saccharolobus solfataricus* (formerly *Sulfolobus solfataricus*) and *Sulfolobus acidocaldarius* cells contain one single genome copy. During replication, three replication origins fire concomitantly once per cell cycle, following a tightly coordinated temporal program reminiscent of the eukaryotic S phase [Lundgren et al., 2004, Robinson et al., 2004, Duggin et al., 2008]. Replication is followed by a defined post-replicative (G2-like) phase before division, highlighting a cell-cycle organization more similar to that of eukaryotes than to bacteria [Lundgren et al., 2004, Bernander, 2007, Gomez-Raya-Vilanova et al., 2025]. By contrast, members of the *Methanobacteriati* kingdom (formerly *Euryarchaea*) possess chromosomes with a single replication origin, with the exception of members of the *Halobacteriota* phylum (formerly *Haloarchaea*) which possess chromosomes with multiple origins of replication. *Halobacteriota* cells contain multiple copies of the genome [Breuert et al., 2006], and DNA replication appears to occur continuously rather than being restricted to a specific cell-cycle window [Zerulla et al., 2014a]. How these highly polyploid organisms regulate the activation and timing of their multiple replication origins in coordination with cell growth and division has yet to be elucidated.

*Haloferax volcanii* is one of the archeaon model organism in which replication has been studied [Greci and Bell, 2020, Perez-Arnaiz et al., 2020]. According to various studies using qPCR, the average number of chromosome copies per cell in exponential phase varies from 12 to 40, and this number decreases as the cells enter stationary phase [Zerulla et al., 2014b, Breuert et al., 2006, Maurer et al., 2018, Ludt and Soppa, 2018]. FACS analyses have also revealed that the ploidy level is highly heterogeneous within the population [Hawkins et al., 2013]. Replication profiling by Marker Frequency Analysis has shown that the laboratory strain H26 carries a 3.5 Mb main chromosome where all four replication origins are active at the population level [Hawkins et al., 2013]. Remarkably, none of them are essential for viability, revealing an alternative mode of replication initiation independent of origins. This origin-independent mode of initiation is dependent upon recombination, indicating that replication initiation can rely on recombination intermediates. Moreover, a strain lacking all four origins shows no significant growth defect, indicating that Recombination-Dependent Replication (RDR) is as efficient as Origin-Dependent Replication (ODR) for growth in optimal laboratory conditions [Hawkins et al., 2013]. Note that since its discovery in *H. volcanii*, Recombination-Dependent Replication has also been reported in *Pyrococcus furiosus*, *Thermococcus kodakarensis* and *Thermococcus barophilus* [Gehring et al., 2017, Mc Teer et al., 2024]. Thus, for a better understanding of both ODR and RDR in *H. volcanii*, we investigated replication dynamics in presence or in absence of the 4 canonical replication origins.

To gain insights into *H. volcanii* replication, we previously implemented optical microscopy techniques and investigated replisome localisation in live cells using widefield microscopy and super-resolution Structured Illumination Microscopy (SIM) [Delpech et al., 2018]. By expressing the single strand-DNA binding protein RPA2 fused with GFP, we provided the first dynamic visualization of DNA replication foci. In this organism, 3 RPA proteins are found but they do not form a heterotrimer [Stroud et al., 2012]. Instead, for RPA1 and RPA3 a unique complex is formed between the RPA and its so-called RPA-associated protein (RPAP) (the *rpa* and the *rpap* genes exist in operons), forming RPA1-RPAP1 and RPA3-RPAP3 complexes. By contrast, RPA2 appears as a monomeric RPA [Stroud et al., 2012]. Of the three *rpa* genes, only *rpa2* is essential for cell viability, highlighting its critical role in DNA replication [Stroud et al., 2012, Skowyra and MacNeill, 2012], and we previously showed that RPA2 is indeed a relevant proxy for localising foci, reflecting active replication sites in the cells [Delpech et al., 2018]. RPA3 is reported to be primarily involved in DNA repair [Stroud et al., 2012]. Strikingly, although no defects in replication or repair are observed in RPA1-deficient cells, the requirement for either RPA1 or RPA3 for viability suggests that RPA1 also contributes to genome maintenance, albeit its exact role remains to be determined.

We revealed that the vast majority of cells contained 1 to 5 RPA2 foci (with an average of 3.3 foci per cell). Surprisingly, this limited number of foci contrasts sharply with the reported high ploidy of *H. volcanii* and the presence of multiple origins per chromosome. In eukaryotes, it has been proposed that such foci may represent the coordinated activity of neighboring replicons and reflect higher-order replication domains rather than individual forks. Whether analogous structures exist in Archaea — and in *H. volcanii* in particular — remains unknown. In order to characterise the replication foci observed in living cells more precisely, we implemented two single molecule localisation microscopy methods: (Fluorescence) Photo-Activated Localisation Microscopy (PALM) on live cells [Betzig et al., 2006, Hess et al., 2006, Turkowyd et al., 2020], and (direct) Stochastic Optical Reconstruction Microscopy (STORM) [Rust et al., 2006, Heilemann et al., 2008] imaging on fixed *H. volcanii* cells using an immunolabelling protocol of RPA2 coupled with a FLAG-tag. Using sptPALM we identified the fraction of mobile and immobile RPA2, and confirmed that the replication foci represent bound RPA2. STORM enabled us to achieve for the first time a sub-40 nm resolution of RPA2 proteins distribution in fixed cells, revealing a significantly higher number of replication foci than previously detected using conventional microscopy. Notably, these replication foci appear to be organised into clusters, a spatial arrangement that could be reminiscent of replication domains described in human cells [Hyrien et al., 2025]. We further showed that lowering the temperature to reduce the growth rate does not affect either the number of replication foci or their spatial organisation at the cellular level. Similarly, altering the mode of replication initiation by inactivating all four replication origins does not disrupt the global replication dynamics, suggesting the existence of a higher-order regulatory layer acting upstream of origin-dependent replication control in *H. volcanii*.

## 2 Results

### 2.1 The number of replication foci increases with cell size

Using widefield fluorescence microscopy, we measured the number of GFP::RPA2 foci and, critically, we improved previous imaging protocols [Delpech et al., 2018] to enable robust cell segmentation from differential interference contrast (DIC) images which allowed us to measure cell surface area. The number of GFP::RPA2 replication foci (RF) detected in exponentially growing cells (OD_600_ *_nm_* ranging from 0.05 to 0.2, generation time *≈*2 h) covers a broad distribution with an average number of 3.56 foci and a standard deviation of 2.04 (n=1140 cells, Figure 1). We previously showed that the number of foci decreases when population growth slows down, but we searched further to identify conditions under which replication foci are no longer observed or are observed very rarely. We observed on average 1.6 (*±*1.2 s.d.) foci per cell in cells transitioning into late exponential phase (OD_600_ *_nm_* ranging from 0.8 to 1, with a generation time of 4 h) which decreased to 0.41 (*±*0.3 s.d.) for cells in the stationary phase (OD_600_ *_nm_ >* 2.0, minimal growth) (Figure 1.b and Supplementary Fig. S1). These observations allowed us to identify specific growth conditions in which replication foci are rarely observed, thereby enabling a direct comparison between cells with active replication and those in which replication has slowed to negligible levels.

**Figure 1:**
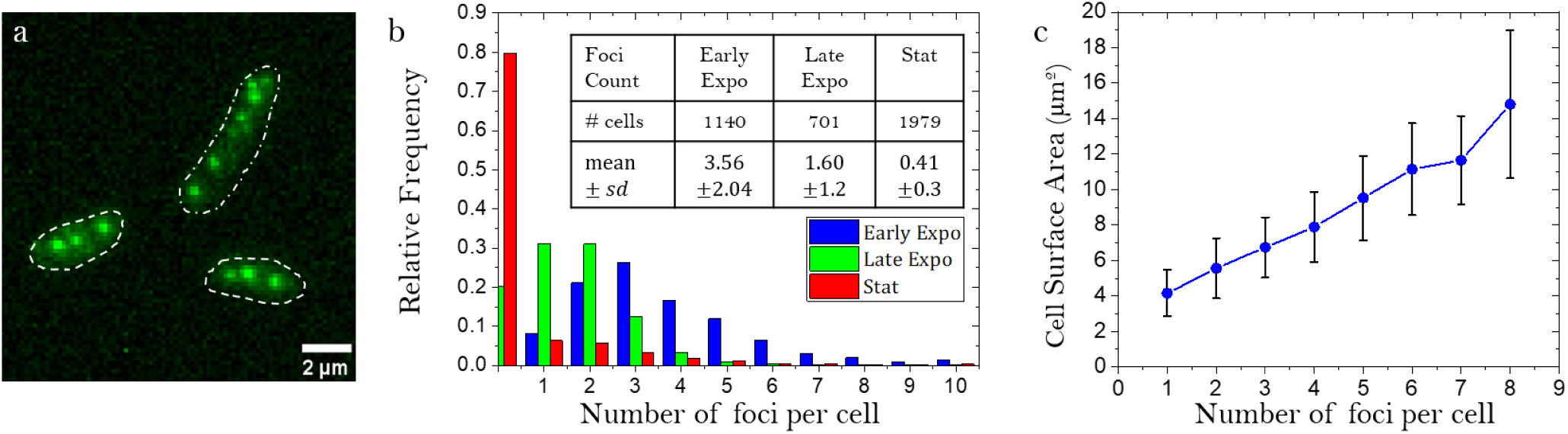
Quantifying the number of replication foci as a function of growth phase and cell size.(a) Example of a widefield image of GFP::RPA2 in exponentially growing cells (b) Distribution of the number of GFP::RPA2 foci per cell in different growth conditions (c) Correlation between the number of foci and the mean cell surface area in exponentially growing cells (error bars represent the standard deviation)

In the asynchronous population of exponentially growing cells, the number of replication foci (RF) ranged from 1 to 8 in the vast majority of cells (Figure 1.b). Strikingly, the fraction of cells devoid of RF remained negligible (Figure 1.b), suggesting there is no cell-cycle phase where replication is absent.

Furthermore, a clear positive correlation was observed between RF number and cell size in exponentially growing cells: Larger cells systematically harbored more replication foci (Figure 1.c). This relationship is consistent with previous reports in *H. volcanii* [Delpech et al., 2018] and *H. salinarum* [Zerulla et al., 2014a], and further supports the notion that replication activity is maintained throughout the cell cycle without an apparent deceleration prior to cell division. Interestingly, this correlation decreases strongly at the end of exponential growth and is absent in stationary cells (Supplementary Fig. S2).

However, our widefield microscopy approach is limited by the Abbe spatial resolution limit of *∼*250 nm, and this correlation between size and number of replication foci could be simply due to the method’s inability to measure a large number of neighbouring foci in a small cell. To overcome this limitation, we used two single molecule super-resolution imaging methods, one applicable to live cells and the other to fixed cells, enabling a more detailed characterization of the observed replication foci.

### 2.2 spt-PALM in live cells identifies bound and unbound RPA2 proteins

Due to this potential bias caused by the resolution limit, we used super-resolution microscopy, starting with photoactivated localisation microscopy (PALM) which improves resolution in living cells [Betzig et al., 2006, Hess et al., 2006]. For this purpose, following [Turkowyd et al., 2020], we engineered a strain expressing the photoactivatable protein Dendra2Hfx fused to the N-terminus of RPA2 from the native chromosomal locus in a background-reduced strain in which the carotenoid synthesis pathway is interrupted (see Methods). Like the GFP::RPA2 fusion protein, Dendra2Hfx::RPA2 appears fully functional. Cells expressing Dendra2Hfx::RPA2 have a growth rate similar to WT cells (Supplementary Fig. S1). Moreover using widefield microscopy, the number of Dendra2Hfx::RPA2 foci is similar to the one observed with GFP::RPA2 (Supplementary Fig. S3).

As shown in Figure 2, PALM imaging of this Dendra2Hfx::RPA2 strain demonstrates a similar distribution of RPA2 replication foci as found in widefield imaging. However, the increased resolution provided by PALM begins to show the additional presence of some sub-structural organization within these foci, hinting that these larger replication foci found in widefield imaging may actually correspond to a more complex cluster of many smaller foci. Unfortunately, the time required to acquire the images was often sufficient to blur these substructures, despite their slow movement, thus reducing the overall improvement in resolution and making an accurate measurement of these substructures difficult.

**Figure 2:**
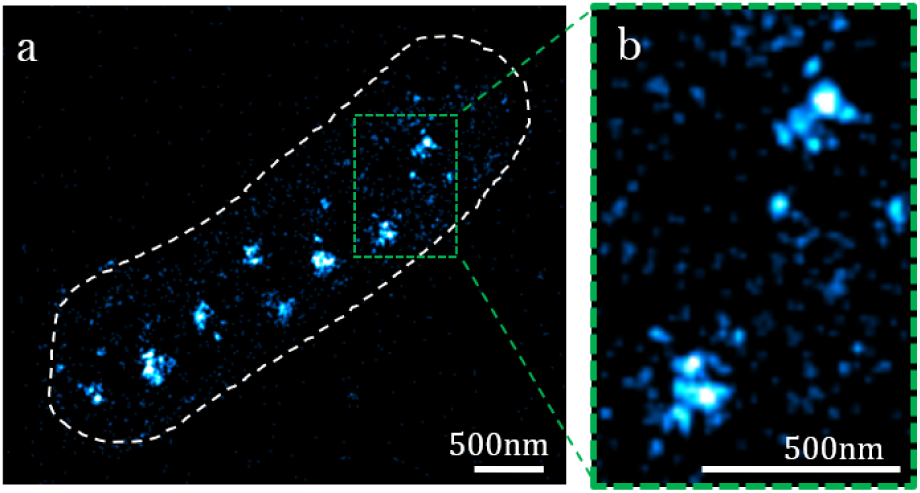
Live PALM imaging of replication foci during early exponential growth. (a) PALM image of Dendra2Hfx::RPA2, with cell outline in white. (b) Zoom on 2 replication foci in the region shown in dashed green, showing some hint of sub-structure (scalebars =500 nm).

To complement PALM imaging, the dynamics of RPA2 proteins in the cells were investigated using single particle tracking PALM (spt-PALM [Manley et al., 2008, Turkowyd et al., 2020]), where the position of single molecules is tracked over several time points to provide information about protein movements. Analysis of the tracks obtained in exponentially growing cells, plotted by their weighted mean jump distance (MJD), revealed two distinct molecular populations: one immobile and the other mobile (Figure 3.a2). By color-coding each track by its classification as immobile or mobile (Figure 3.a1), we could observe that immobile molecules are localised primarily in foci, while mobile molecules are present everywhere in the cell. These results are consistent with the observed fluorescent foci being indeed replication foci formed by RPA2 bound to single-stranded DNA and therefore immobile, while unbound RPA2 molecules are moving throughout the cell and contribute a diffuse background signal. Similar spt-PALM experiments in stationary cells showed both a decrease in the number of tracks (Supplementary Fig. S4) as well as a decrease in the ratio between bound and mobile RPA2 proteins (Figure 3.d). This indicates that RPA2 expression level is dependent on growth phase but that the stronger decrease in replication activity in stationary phase results in an excess of RPA2 compared to the number of replication forks.

**Figure 3:**
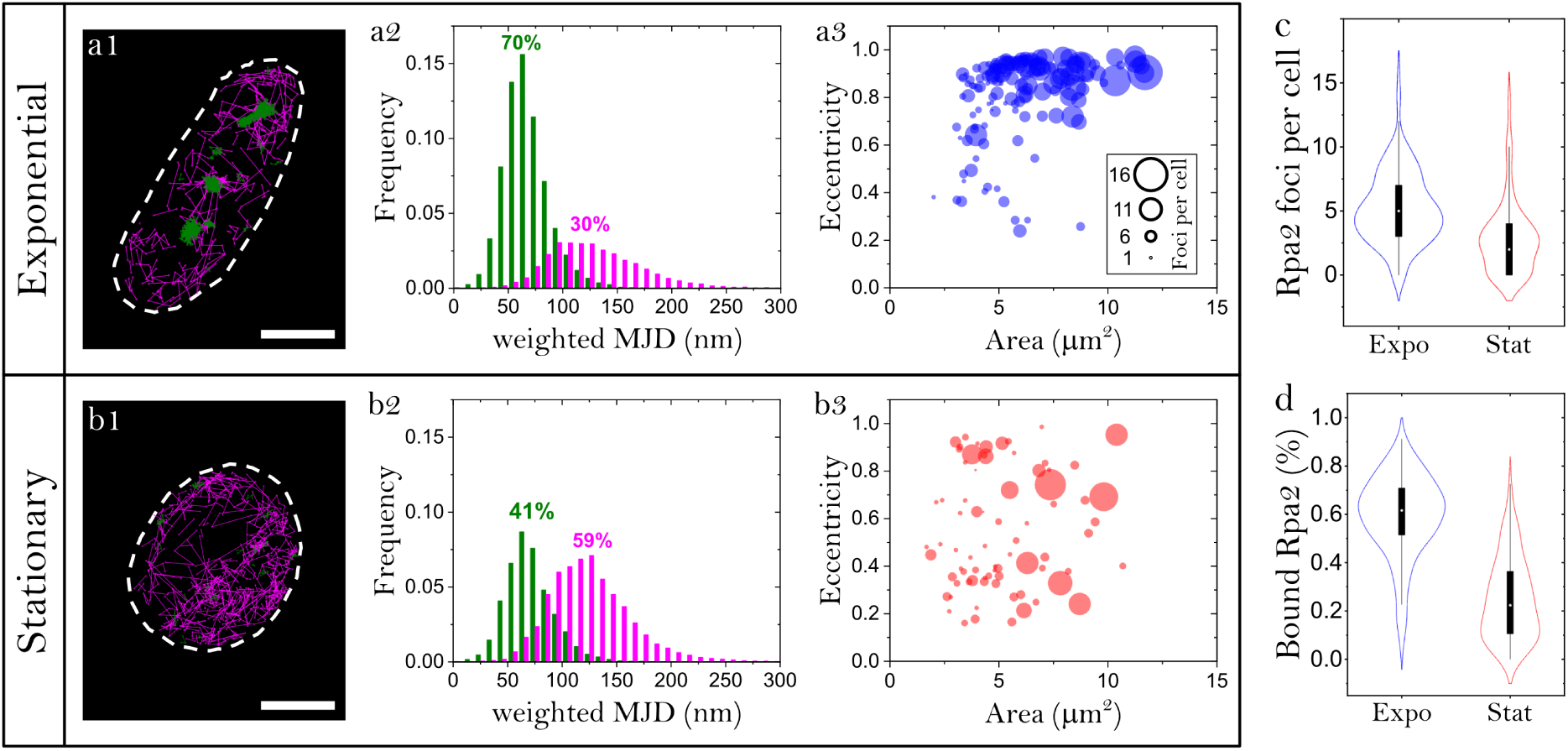
Single-particle tracking PALM image of Dendra2Hfx::RPA2 *H. volcanii* shows foci formed by immobile RPA2, assumed to be bound to single-stranded DNA, and mobile RPA2 everywhere in the cell with differences in cell shape, foci number and size between exponential (a1) and stationary (b1) cells (scalebar =1 *µ*m). Relative frequency of weighted mean jump distances for immobile (green) and mobile (pink) Dendra2Hfx::RPA2 tracks in exponential (a2) and stationary (b2) *H. volcanii* cells. Number of foci per cell formed by immobile localisations in exponential (a3) and stationary (b3) phase with respect to cell area in *µm*^2^ and cell eccentricity. Number of foci and area are correlated in both growth phases (Expo: Pearson Corr.=0.54, p*<* 0.0001; Stat: Pearson Corr.=0.43, p*<* 0.0001). Correlation between eccentricity and number of foci is less significant (Expo: Pearson Corr.=0.29, p=1.9E-4; Stat: Pearson Corr.=0.017, p=0.86 not significant) (c) Number of foci per cell formed by immobile RPA2 (Mann-Whitney test, p*<* 0.0001). (d) Percentage of bound RPA2 localisations (Mann-Whitney test, p*<* 0.0001).

We then quantified the number of foci formed by localisations classified as immobile. Consistent with the widefield measurements (Figure 1.b), the number of foci per cell decreased from exponential to stationary phase (Figure 3.c).

Comparing the number of foci with cell morphology confirmed that cell size was the strongest predictor of foci number in exponentially growing cells (Figure 3.a3). As a second parameter, we examined cell eccentricity, which has been established as a marker of rapid growth in *H. volcanii* [de Silva et al., 2021]. Although eccentricity showed a weaker correlation with foci number than cell size, it revealed the heterogeneity within exponentially growing cultures. Most cells displayed high eccentricity, indicative of rapid growth, with larger cells containing more RPA2 foci. In contrast, a subpopulation of smaller, rounder cells exhibited fewer RPA2 foci, consistent with reduced growth (Figure 3.a3). These observations indicate that, despite the overall ensemble characteristics of exponential cultures, individual cells span a range of growth states. Conversely, stationary-phase cultures still contained occasional larger, more elongated cells, but the relationships between cell size, eccentricity, and foci number were largely lost (Figure 3.b3). This is consistent with the increased physiological heterogeneity of stationary-phase populations, in which individual cells can occupy different metabolic and growth states despite exhibiting similar overall morphology. As a result, cell morphology becomes a much poorer predictor of RPA2 focus formation than during exponential growth.

Taken together, increasing the resolution by using PALM and spt-PALM imaging provided a more detailed view of the replication foci and confirmed the results obtained with widefield imaging. Moreover, the data hints at a sub-organisation in the replication foci. To determine their organisation at the nanoscale, we then turned to structural, fixed cell STORM super-resolution imaging.

### 2.3 STORM super-resolution on fixed cells can resolve the nano-organization of replication foci

To investigate possible substructures in the replication foci observed in widefield and PALM with higher spatial resolution, we turned to fixed cells with immunolabeling to perform STORM microscopy [Rust et al., 2006, Heilemann et al., 2008]. We engineered a fully functional strain expressing FLAG::RPA2 from its native chromosomal locus (Supplementary Fig. S1). We then performed immuno-labeling, enabling specific and efficient detection of RPA2. This strategy allowed us to establish the first STORM-based super-resolution imaging approach of a protein in *Haloferax volcanii*. STORM images (Figure 4) uncovered clear substructures within replication foci that appeared as single diffraction-limited spots in wide-field imaging (Figure 4.a1,a2) resulting in a substantially larger number of detectable RPA2 foci (Figure 4.d).

**Figure 4:**
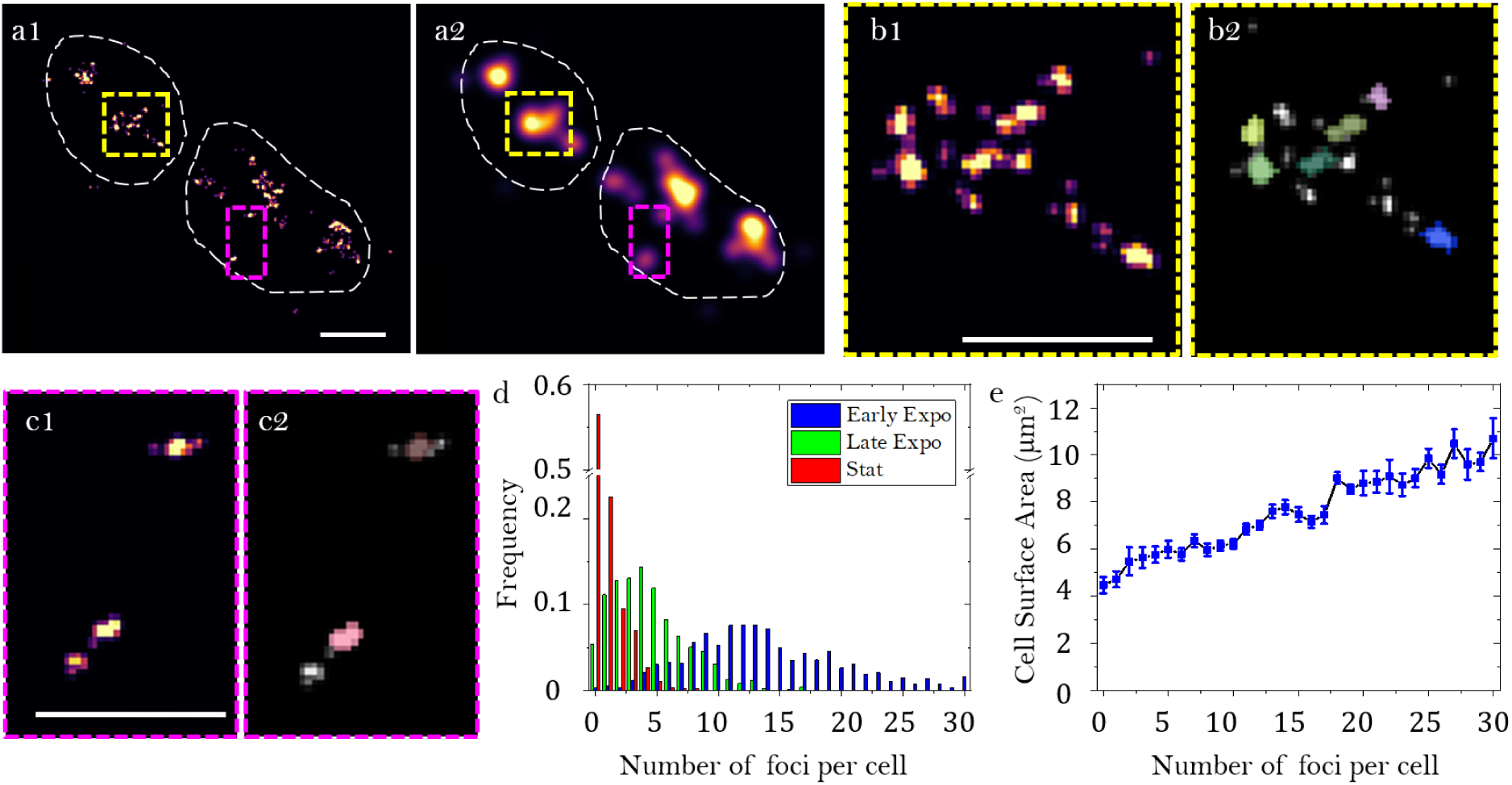
(a1) STORM image of RPA2 proteins in *Haloferax volcanii* cells in early exponential growth (cells outlined in yellow) and (a2) the corresponding reconstructed widefield image. (b1) Zoom onto the yellow zone highlighted in (a1), with (b1) the raw data and (b2) the segmented data. (c1) Zoom onto the purple zone highlighted in (a1), with (c1) the raw data and (c2) the segmented data. (d) Distribution of the number of FLAG::RPA2 foci per cells for the 3 different growth conditions chosen: early exponential in blue (n=1275), late exponential in green (n=733) and stationary in red (n=1233). (e) Average cell surface area for the cells that have exactly *n* foci in exponentially growing cells, with *n* between 0 and 30. (scalebar =1 *µ*m in a1,b1 and c1. Error bars represent the standard deviation)

We quantified a mean number of FLAG::RPA2 foci per cell of 13.75 *±* 0.35 in exponential phase (OD_600_ *_nm_* ranging from 0.05 to 0.2; *n* = 1,275 cells), compared to 0.057 *±* 0.03 in the negative-control strain lacking the FLAG tag (*n* = 684 cells) which means false-positive detections represent less than 0.5% of the foci detected in FLAG::RPA2 cells. Consistent with our widefield observations, we found a marked decrease in the number of FLAG::RPA2 foci as cells transitioned to stationary phase: The average number dropped to 4.49 *±* 0.21 during late exponential growth (OD_600_ *_nm_* ranging from 0.8 to 2.0; *n* = 733 cells) and further declined to 0.84 *±* 0.20 in stationary phase (OD_600_ *_nm_ >* 2.0, *n* = 1,233 cells) (Figure 4.d) indicating a reduction in replication activity concomitant with the slowdown of cellular growth.

In exponentially-growing cells, the number of FLAG::RPA2 foci was highly heterogeneous, ranging from few foci up to 30 or more foci (Figure 4.d). Notably, the fraction of cells lacking foci remained very low (0.7 *±* 0.9%). This observation further confirmed the absence of a strict cell-cycle specific control and the lack of a gap period prior to and/or after cell division. Importantly, the correlation between cell size and number of replication foci observed in exponentially growing cells by widefield imaging of GFP::RPA2 foci holds when using STORM to quantify the number of FLAG::RPA2 foci (Figure 4.e). Furthermore, a clear correlation could also be observed between foci number and DNA content quantified using Hoechst: the higher the DNA content, the higher the number of foci (Supplementary Fig. S5). We also observed a positive correlation between cell size and DNA content with larger cells having higher DNA content (Supplementary Fig. S5). As in the case of widefield imaging (Supplementary Fig. S2), this correlation weakens substantially at the end of exponential growth, and is completely lost in the stationary phase (Supplementary Fig. S5). Taken together, these results suggest that in exponentially growing cells the DNA content increases with cell size and higher DNA content correlates with increased replication activity, but this correlation is lost as cells enter the stationary phase.

### 2.4 STORM reveals clustered organization of replication foci

Strikingly, FLAG::RPA2 foci are not evenly distributed throughout the cell; rather, regions with a higher density of foci could be observed. Indeed, displaying the distance to the nearest neighbour of all foci in the population of cell (Figure 5a) shows a probability of 75% of foci having at least one neighbour within 250 nm, which corresponds to the size of the replication foci observed with widefield microscopy. To further investigate the spatial distribution of replication foci we used the clustering algorithm Density-based spatial clustering of applications with noise (DBSCAN) [Ester et al., 1996, Endesfelder et al., 2013]. We distinguished isolated replication foci from clustered ones by setting in DBSCAN a maximum distance (*ɛ*) of 250 nm and a minimum number of points required to form a cluster (minPts) equal to 2 (Figure 5b). No significant difference in signal intensity was detected between clustered and isolated replication foci (24, 4 *±* 1, 7 and 24, 0 *±* 1, 3 respectively), indicating that individual foci contain comparable amounts of FLAG::RPA2.

**Figure 5:**
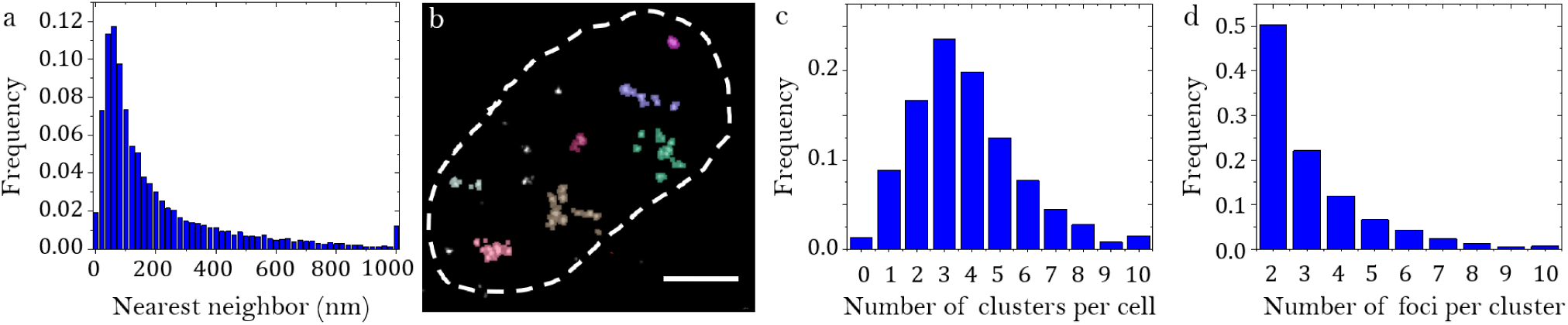
Organisation of clusters of FLAG::RPA2 foci in early exponentially growing cells. (a) Distance to the nearest foci calculated in WT exponentially growing cells, (b) Example of a cell in which the DBSCAN clustering algorithm was applied, which groups neighbouring foci together into clusters, displayed here by color, with the cell outline in dashed line. (c) Number of clusters detected per cell in exponentially growing cells. (d) Number of foci per cluster in exponentially growing cells

In exponentially-growing cells, isolated RF represented 36.4 *±* 1.6% of RF per cell on average, with a decrease in the proportion of isolated RF as the total number of RF increases. As for the identified clusters of RF, they contain on average 3.21 *±* 0.07 RF (Figure 5.b). The number of clusters ranged from 1 to 8 in the vast majority of cells, reminiscent of the widefield quantification (Figure 1.d). By focusing on clusters containing at least three foci, we measured an average distance between foci of 134.7 *±* 1.1 nm.

Overall, these observations show that most RPA2 foci cluster into denser regions corresponding to the replication foci detected in wide-field imaging, whereas isolated RPA2 foci are only resolved by STORM. We then tested whether this organisation is specific to the exponentially growing phase in wild type *Haloferax volcanii* in optimum growth conditions, first by changing the temperature.

### 2.5 Temperature-dependent modulation of the replication program

Environmental changes such as variations in nutrient availability, temperature, salinity, and pH trigger cellular stress and adaptation mechanisms that can influence the DNA replication program. As previously observed when comparing exponential and stationary phases, increased cell density and nutrient limitation reduce both the growth rate and the frequency of replication initiation. To further investigate this regulatory response, we examined the impact of a moderate temperature shift on replication dynamics. *Haloferax volcanii* was cultured at 37°C—a suboptimal temperature in rich medium—where it exhibited a 50% increase in generation time (188 min) compared with growth at 45°C (118 min) (Supplementary Fig. S1).

Using STORM microscopy, we noted a significant decrease in the number of replication foci observed in exponentially growing cells at 37°C compared to 45°C. The mean number of FLAG::RPA2 foci per cell was 8.03 *±* 0.41 at 37°C (*n* = 1,275 cells), compared to 13.75 *±* 0.35 at 45°C (*n* = 684 cells) (Figure 6.a). We also observed a decrease in cell size, with an average size of 6.12 *µm*^2^ *±* 0.17 at 37°C compared to 7.40 *µm*^2^ *±* 0.40 at 45°C (Figure 6.b). However, the same correlations were observed: the larger the cells, the greater their DNA content (Supplementary Fig. S5) and the greater their number of replication foci (Figure 6.c). These results suggest that slowing down growth limits the maximum cell size, DNA content, and number of replication foci of cells within the population. Nevertheless, an equal number of replication foci indicates similar size and DNA content. Similarly, the spatial organisation of replication foci at 37°C is identical to that observed at 45°C: The proportion of clustered foci, the number of replication foci per cluster, and the signal intensity for both isolated and clustered foci are all similar (Supplementary Fig. S5). These results all suggest that the cells at 37°C mirror a subpopulation of small cells at 45°C with similar replication dynamics.

**Figure 6:**
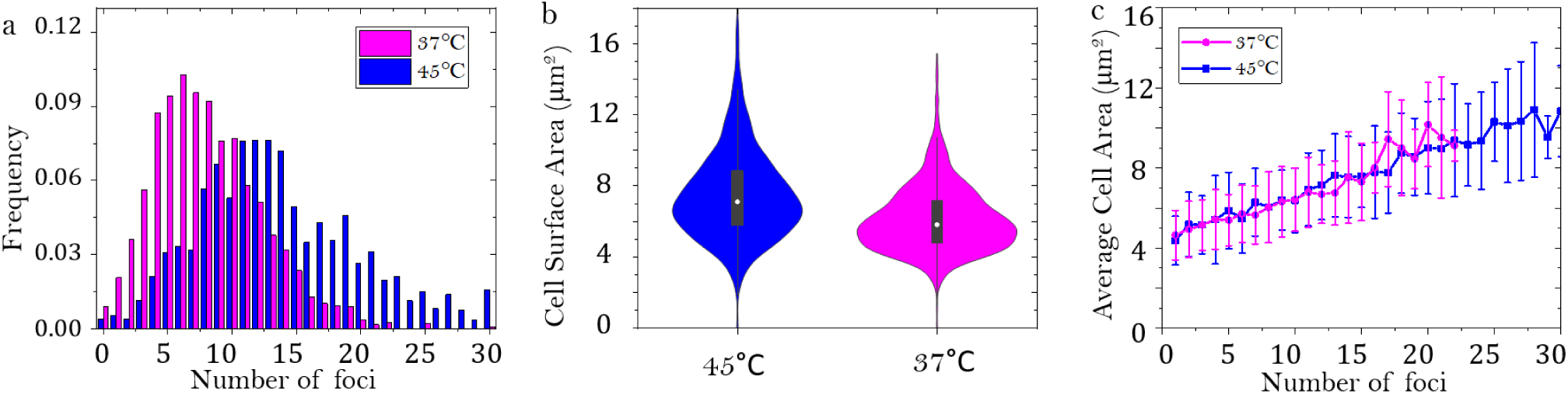
Replication dynamics at 37°C. (a) Distribution of detected number of foci per cell using STORM microscopy in exponentially growing cells at 45°C (blue) versus 37°C (magenta). (b) Corresponding size distribution of the population. (c) Average cell size at 37°C and 45°C for a given number of foci detected per cell. (error bars represent the standard deviation)

### 2.6 Replication foci organization does not rely on origin firing

We then investigated whether the alternative mode of replication initiation relying on homologous recombination (Recombination-Dependent Replication) could affect the organisation of replication. Δ*ori* cells lacking all four replication origins exhibit faster growth than wild-type (WT) cells in competitive growth assays, despite maintaining similar DNA content [Hawkins et al., 2013]. We examined the spatial organisation of replication foci in such strain using STORM microscopy. For that purpose, the *rpa2* gene was replaced by its *flag::rpa2* version at the chromosomal locus in a Δ*ori* background [Hawkins et al., 2013]. The FLAG-tagged version of RPA2 is fully functional in this background, as *flag::rpa2* Δ*ori* cells exhibit a growth rate comparable to that of their *flag::rpa2* counterparts (Supplementary Fig. S1).

We quantified the mean number of FLAG::RPA2 foci per cell as 9.77 *±* 0.49 in exponential phase (OD_600_ *_nm_* from 0.05 to 0.2; *n* = 306 cells) (Figure 7.a). As observed in the presence of the four replication origins, when the cells transitioned to the stationary phase, there was a marked decrease: The average number of foci dropped to 3.49 *±* 0.32 upon entry into the stationary phase (OD_600_ *_nm_* ranging from 0.8 to 2.0; *n* = 477 cells) and declined further to 0.99 *±* 0.11 in the late stationary phase (OD_600_ *_nm_ >* 2.0, *n* = 689 cells) (Supplementary Fig. S7).

**Figure 7:**
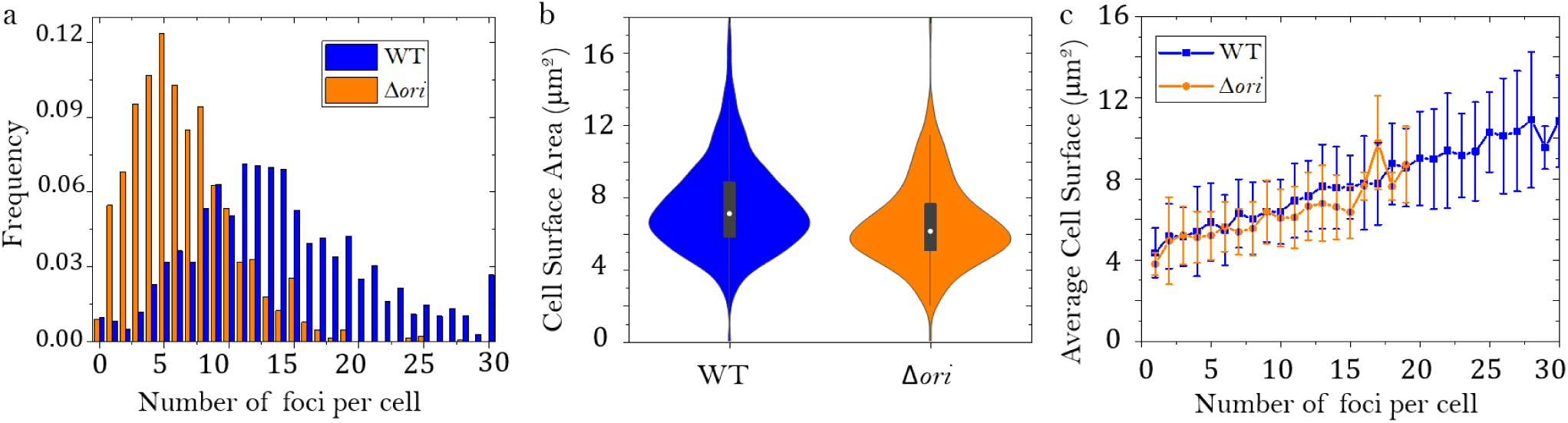
Replication dynamics in Δ*ori* mutant (a) Distribution of detected number of foci per cell using STORM microscopy in exponentially growing cells at 45°C in the wildtype (blue) and Δ*ori* mutant (orange). (b) Corresponding size distribution of the population. (c) Average cell size in the wildtype (blue) and Δ*ori* mutant for a given number of foci detected per cell (error bars represent the standard deviation).

We observed a decrease in cell size (Figure 7.b), with an average size of 6.56 *µm*^2^ *±* 0.18 compared to 7.40 *µm*^2^ *±* 0.40 in the wild type. However, the same correlations were observed: the larger the cells, the greater their DNA content and the greater their number of replication foci (Figure 7.c). We noticed that exponentially growing Δ*ori* cells at 45°C show a fairly similar distribution of number of replication foci to exponentially growing WT cells at 37°C. Yet, growing Δ*ori* cells at 37°C induced a decrease in the number of FLAG::RPA2 foci per cell to 6.57 *±* 0.50 (*n* = 862 cells) while the average size remained unchanged with 6.17 *µm*^2^ *±* 0.26. Although the same correlations were observed — that is, the larger the cells, the greater their DNA content and number of replication foci — these correlations are somewhat weaker (Supplementary Fig. S8).

More surprisingly, the spatial organisation of replication foci in the Δ*ori* context is identical to that observed in the WT context. The presence and proportion of clustered foci, the number of replication foci per cluster, and the signal intensity for both isolated and clustered foci are all highly similar (Supplementary Fig. S7). Moreover, when comparing the spatial organisation of replication foci at 37°C between WT cells and Δ*ori* cells, there is an increase in the proportion of isolated foci in Δ*ori* cells reflecting a weaker organisation of replication foci into clusters.

Thus, the absence of replication origins does not affect the number of replication foci observed or their spatial organisation at 45°C for a given cell size (Supplementary Fig. S8). However, relying on RDR for replication limits the maximum cell size, DNA content, and number of replication foci of cells within the population.

## 3 Discussion

To localise replication sites in *Haloferax volcanii*, we targeted the key replication protein RPA2. In exponentially-growing cells, replication foci formed by GFP::RPA2 proteins can be observed using widefield fluorescence microscopy [Delpech et al., 2018]. However, these observations are limited by the *∼*250 nm resolution of the imaging technique. To overcome these limitations and better characterize replication sites in *Haloferax volcanii*, we implemented two single molecule super-resolution modalities: (spt-)PALM and STORM. Using spt-PALM we could confirm that the replication foci identified consist of immobile RPA2 proteins, most likely bound to DNA, and that a second population of mobile proteins is present in the cell. This method also gave a glimpse of sub-structure in the replication foci, which we further studied using immunolabeling of a FLAG tagged version of RPA2 coupled to STORM imaging. STORM imaging of FLAG::RPA2 foci at *∼*30 nm resolution revealed that live foci corresponded to multiple replication foci too close to be resolved at 250 nm resolution, and allowed the observation of isolated replication foci that could not be detected otherwise. Our data provides a better understanding of how replication is coupled to the cell cycle during exponential growth. The size of cells within the exponentially-growing population shows significant variability, exceeding a factor of two. Time-lapse study of the archaeon *Halobacterium salinarum* has demonstrated that it controls its size by adding a constant length between two cell cycle events, similar to bacteria and fungi, but with higher variance in cell division ratio and growth rate, which explains the greater size heterogeneity found in the population [Eun et al., 2018]. Our data suggest that the same relaxed cell cycle control mechanisms could apply to *H. volcanii* cells. Furthermore, our results demonstrate that there is no cell cycle phase where replication is absent, as suggested for *H. salinarum* [Zerulla et al., 2014a]. Indeed, a cell cycle control inhibiting replication before and after division would result in the presence of a population of small and large cells within the population (i.e. after and prior to cell division, respectively) without replication foci. But cells in which no replication foci can be detected only correspond to the smallest cells, and represent a very limited proportion of cells (0.7%). Moreover, we show that an increase in size is associated with an increase in DNA content. These observations suggest that, at the level of the individual cell, an increase in size during the cell cycle is accompanied by a proportional increase in the number of genome copies such that a constant proportion of chromosomes are replicated. Consistently, an increase in size and DNA content is associated with an increase in the number of replication foci. Overall, our results suggest that the cell cycle mechanisms that enable a constant increase in size allow for a coupled increase in the number of chromosome copies through constant replication rate during the cell cycle. It may be mediated by the dose-dependent activity of key limiting factors, such as replication initiators and/or metabolic regulators. The absence of a fixed upper limit suggests that these factors are not subject to strict stoichiometric constraints. Instead, they react dynamically, with their abundance increasing in proportion to cell size and/or chromosome copy number (provided that each copy contributes equally to gene expression). It remains to be determined what proportion of chromosomes are replicated in each cell of *H. volcanii*, and what are the molecular mechanisms of cell cycle control. Polyploid bacteria, such as cyanobacteria, represent additional systems in which chromosome replication occurs in a multi-copy genomic context [Ohbayashi et al., 2019]. However, in contrast to monoploid bacterial models—where replication dynamics have been extensively characterized [Willis and Huang, 2017]—these processes remain poorly understood in polyploid organisms, largely due to the lack of single-cell microscopy data. Together, these systems provide a relevant comparative framework, in which studying *H. volcanii* could yield critical insights into the general principles, as well as the specific challenges and adaptations, of DNA replication in polyploid cells.

Remarkably, this control of the cell cycle allowing a coupled increase of cell size and chromosome copies is preserved in conditions of slower growth. Indeed, we show that a 50% increase in the doubling time of *H. volcanii* cells by growing them at 37°C does not significantly impact this homeostasis. The slower growth rate limits the maximum cell size, DNA content, and number of replication foci of cells within the population. However, if we compare the two cell populations with an equivalent number of replication foci, the size of the cells is comparable. Assuming that similar sizes indicate similar numbers of chromosome copies and that the replication program still relies on the firing of the four canonical replication origins, these observations suggest that slow growth also induces a decrease in replication speed, as has been reported in bacteria [Bhat et al., 2022]. Nonetheless, a thorough assessment of how slow growth affects ploidy levels and replication rate is required for robust conclusions.

More surprisingly, our results show that cells relying on Recombination-Dependent initiation of replication (RDR) retains a similar control of the cell cycle. Δ*ori* cells show similar growth rate when compared to wild-type cells. Nevertheless, we show that in Δ*ori* cells the maximum cell size, DNA content, and number of replication foci is lower than in WT cells. However, if we compare the two cell populations with an equivalent number of replication foci, the average size of the cells is once again comparable. As the DNA content of Δ*ori* cells was previously reported to be similar to that of WT cells [Hawkins et al., 2013], these results suggest that control of the cell cycle, which enables synchronised increased in cell size and in the number of chromosomes, is maintained even though precise regulation of replication initiation at origins is abolished.

Thus, our results argue against a single cell cycle control occurring at the initiation of DNA replication, as proposed for *E. coli* [Zheng et al., 2016], leaving open the question of the underlying molecular mechanisms responsible for the cell cycle control in both WT and Δ*ori* cells. In this context, which molecular or physiological factors could impose this limitation on cell size and chromosome copies in Δ*ori* cells? Unregulated replication would lead to excessive consumption of energy and/or DNA synthesis precursors, thereby limiting growth. Although we do not observe any clear deregulation in the number of replication foci, this does not rule out the possibility that a greater number of replication forks are present on each chromosome during the cell cycle: Random initiation from recombination intermediates along the chromosome could result in adjacent forks becoming indistinguishable, and could prove to be more metabolically costly than replication control at canonical replication origins.

Secondly, our findings reveal a previously undetected spatial arrangement of replication foci with a majority (80%) forming clusters of 2-4 replication foci. Our findings reveal a two-tiered organization: (i) Individual replication foci, resolved by super-resolution imaging, and (ii) Higher-order replication domains, formed by the spatial grouping of these foci. Our data suggest that individual foci, either clustered or isolated could correspond to individual replication forks. Thus, replication domains are likely revealing a spatial grouping of multiple replication forks, which raises new questions about the organisation and dynamics of replication in *H. volcanii*.

The replication foci previously observed via widefield imaging of GFP::RPA2 can now be interpreted as spatially defined replication domains, each comprising multiple active replication sites. The underlying mechanisms of replication domain formation remain to be understood. One hypothesis is that these replication domains could arise from topological constraints, where DNA segments are physically brought into proximity to facilitate coordinated replication. They could connect multiple replication forks on separate copies of the chromosome, as well as multiple replication forks replicating the same copy. Such clustering of replication forks has been reported in mammalian cells, with the observation of Replication Domains (RDs) corresponding to co-replicating forks starting replication synchronously [Xiang et al., 2018]. This could involve the combined action of the SMC (Structural Maintenance of Chromosomes) protein and transcription, both shown to be driving forces of chromosome structuring in *H. volcanii* [Cockram et al., 2021]. Strikingly, this spatial organization is also conserved in Δ*ori* cells relying on Recombination-Dependant Replication, and the conservation of these features in Δ*ori* cells challenges the hypothesis that the underlying replication regulation mechanism relies on initiation at defined origins. Our work paves the way for a better understanding of the molecular nature of these replication domains in *H. volcanii* and the mechanisms responsible for their formation. Instead, our data support a model where spatial cues or global chromosomal architecture—rather than origin firing—may coordinate replication timing and cell cycle.

Moreover, a much larger number of replication foci could be detected using super-resolution imaging of FLAG::RPA2 than widefield imaging of GFP::RPA2. In exponentially growing cells, we could detect an average of 14 replication foci, with the majority of cells having between 8 and 20 replication foci. What we can learn about the dynamics of replication from the number of foci per cell is a complex question, as the number of replication foci observed at any given time in a cell depends on multiple parameters, including: the number of chromosome copies, the proportion of copies being replicated, the replication program (i.e. how many replication origins are activated per chromosome and when they fire), and the replication fork speed. Furthermore, the technical biases of imaging must also be considered, with the main bias being an underestimation of the numbers if not all are detected, while others could be too close to be resolved.

Therefore, a better understanding of replication dynamics at the cellular scale will require the development of techniques such as Fluorescence *In Situ* Hybridisation (FISH), which could enable an accurate quantification of the ploidy in each cell, or improved protocols for neosynthesized DNA imaging. Indeed, although we have shown that BrdU can be effectively incorporated into DNA in *Haloferax volcanii* [Delpech et al., 2018, Lestini et al., 2022], its specific detection by immunolabelling as *bona fide* replication foci proved too unreliable for short periods of incorporation (less than 15 minutes). Nevertheless, we show that such organisation of replication foci into replication domains is conserved in slower growth conditions, which could be easier to study experimentally.

## 4 Conclusion

Taken together, our data uncover a higher-order spatial organisation of replication where foci cluster into discrete domains containing a few active sites. Furthermore, our findings suggest a coordinated regulation of replication and cell cycle progression, despite the absence of temporal restrictions on replication between divisions. The conservation of spatial replication organisation and cell cycle regulation, even during slow growth or in the absence of all canonical origins, suggests that these processes are governed by robust mechanisms that are independent of the replication initiation mode. These findings redefine our understanding of replication control, revealing that spatial architecture and chromosomal topology—rather than origin-dependent initiation—serve as the primary determinants of replication coordination and cell cycle progression in *Haloferax volcanii*. It would be interesting to establish whether this is a characteristic that is shared by most organisms of the *Methanobacteriati* kingdom or if it is only found in Archaea where replication initiation can rely on either origin firing or recombination (i.e. also in *Pyrococcus furiosus*, *Thermococcus kodakarensis* and *Thermococcus barophilus*).

## A Methods

### A.1 Cell Culture and strain

*Escherichia coli* strains XL1-Blue MRF’ (Δ*mcrA183* Δ*mcrCB-hsdSMR-mrr173 endA1 supE44 thi-1 recA1 gyrA96 relA1 lac [F’ proAB lacIqZ*Δ*M15 Tn10]*) and GM121 (F-*dam-3 dcm-6 ara-14 fhuA31 galK2 galT22 hdsR3 lacY1 leu-6 thi-1 thr-1 tsx-78*) were used for cloning. The latter *dam-dcm-* strain was also used to prepare unmethylated plasmid DNA for efficient transformation of *H. volcanii*. *E. coli* transformants were selected on LB plates containing 100 µg/mL ampicillin, 0.5 mM IPTG and 80 µg/mL X-Gal.

*H. volcanii* cultures using enriched Hv-YPC or Hv-Ca media were grown at 45°C, as described previously [Delpech et al., 2018]. Hv-YPC contained (per liter) 144 g of NaCl, 21 g of MgSO4.7H2O, 18 g of MgCl2.6H2O, 4.2 g of KCl, and 12 mM Tris HCl (pH 7.5), Yeast extract (0.5%, wt/vol; Difco), 0.1% (wt/vol) peptone (Oxoid), and 0.1% (wt/vol) Casamino Acids (Difco). Casamino Acids medium (Hv-Ca) was similar to Hv-YPC except that yeast extract and peptone were omitted, and Casamino Acids were added to a final concentration of 0.5% (wt/vol). 50 µg/mL tryptophan was added to enable the growth of Δ*trpA* auxotrophic strains. The *H. volcanii* strains used in this study and characteristic features are listed in Supplementary Table S1.

#### A.1.1 Construction of the *Haloferax volcanii* strains expressing FLAG::RPA2 and Dendra2Hfx::RPA2

All strains, plasmids and primers are listed in Supplementary Table S1, S2 and S3.

##### FLAG::RPA2 construct

To generate the FLAG::RPA2 construct, the upstream region of the *rpa2* gene (US) and the first 400 bp of the *rpa2* gene (*rpa2’*) were cloned into pTA131 [Allers et al., 2004], with the addition of the FLAG-coding sequence 5’-GACTACAAAGACGATGACGACAAG-3’ at the 5’ end of the *rpa2* gene to generate the pRL83 plasmid. The US region was generated by PCR on H26 genomic DNA using primers RL272 and RL273. The *rpa2’* region was generated by PCR on H26 genomic DNA using primers RL274 and RL275. Each PCR product contained 30 bp homology with adjacent fragments for SLIC cloning [Li and Elledge, 2007]: RL272 shares 30 bp homology with the EcoRI-digested extremity of pTA131, RL273 shares 30 bp homology with the PCR product rpa2’, including the FLAG sequence, RL274 shares 30 bp homology with the US PCR product, including the FLAG sequence, and RL275 shares 30 bp homology with the NotI-digested extremity of pTA131. Following the SLIC method, PCR fragments and the linearised plasmid were digested using T4 DNA polymerase for 45 minutes at 22°C to generate 3’-single-stranded extremities, then all amplification products were mixed in a 1:1:1 molar ratio and incubated for 30 minutes at 37°C before transformation into *E. coli* XL1-blue. Transformants were selected on LB plates containing 100 µg/mL ampicillin, 0.5 µM IPTG, and 80 µg/mL X-Gal. The presence of the correct insert in the plasmid, as determined by white colonies, was tested by colony PCR using primers pBSF2 and pBSR3. The sequence of one selected plasmid, dubbed pRL83, was further confirmed by Sanger sequencing. Then pRL83 was used to transform H26 strain using the pop-in/pop-out method as described previously [Bitan-Banin et al., 2003]. Pop-out colonies were plated on Hv-Ca plates containing 5-FOA. The presence of the FLAG sequence was tested by PCR using primers RL129 and RL287. The *flag::rpa2* sequence was present in 7 colonies out of 10 tested. One FLAG::RPA2 construct was conserved for further studies and dubbed HvRL133. The same method was used to transformed H1804 with pRL83, adding 50 µg/mL of tryptophan to the media for growth. The FLAG::RPA2 sequence was present in 3 colonies out of 8 tested. One FLAG::RPA2 construct was conserved for further studies and dubbed HvRL136.

##### Dendra2Hfx::RPA2 construct

The Dendra2Hfx::RPA2 construct was generated using the same strategy. The upstream region (US), the *dendra2hfx* gene and the first 400 bp of the *rpa2* gene (*rpa2’*) were cloned into pTA131 to generate the pRL118 plasmid. The US region was generated by PCR on H26 genomic DNA using primers RL272 and RL397. The *dendra2Hfx* gene was generated by PCR on the plasmid DNA pTA962-FtsZ1-Dendra2Hfx [Turkowyd et al., 2020] using primers RL398 and RL399. The *rpa2’* region was generated by PCR on H26 genomic DNA using primers RL400 and RL275. Each PCR product contained 30 bp homology with adjacent fragments for SLIC cloning [Li and Elledge, 2007]: RL272 shares 30 bp homology with the EcoRI-digested extremity of pTA131, RL398 shares 30 bp homology with the US product, RL400 shares 30 bp homology with the *dendra2Hfx* product, and RL275 shares 30 pb homology with the NotI-digested extremity of pTA131. Following the SLIC method, PCR fragments and the linearised plasmid were digested using T4 DNA polymerase for 45 minutes at 22°C to generate 3’-single-stranded extremities, then all amplification products were mixed in a 1:1:1:1 molar ratio and incubated for 30 minutes at 37°C before transformation into *E. coli* XL1-blue. Transformants were selected on LB plates containing 100 µg/mL ampicillin, 0.5 µM IPTG, and 80 µg/mL X-Gal. The presence of the correct insert in the plasmid, as determined by white colonies, was tested by colony PCR using primers pBSF2 and pBSR3. The sequence of one selected plasmid, dubbed pRL118, was further confirmed by Sanger sequencing. Then pRL118 was used to transform Δ HVO 2528 strain given by Jörg Soppa using the pop-in/pop-out method as described previously [Bitan-Banin et al., 2003]. Pop-out colonies were plated on Hv-Ca plates containing 5-FOA. The presence of *dendra2Hfx* sequence was tested by PCR using primers RL401 and RL403. 2 colony out of 15 contained the *dendra2Hfx* gene, one was conserved for further studies and dubbed HvRL217. The chromosomal insertion of *dendra2Hfx* sequence at RPA2 locus was further confirmed by PCR on HvRL217 genomic DNA using primers RL141 and RL165.

#### A.1.2 Cell preparation for STORM microscopy

Once the cell culture has reached the targeted OD, 2% formaldehyde is added to the culture for 20 minutes, followed by the addition of Glycine at a final concentration of 110 mM for 5 minutes. Formaldehyde and Glycine incubation were done at 45°C with agitation at 150 rpm, as for cell growth. Next steps were all performed at room temperature. Cells were harvested by centrifugation for 8 minutes at 2,410 RCF (Relative Centrifugal Force, or g force). Cells were washed with 1X PBS and then harvested by centrifugation for 8 minutes at 2,410 RCF and re-suspended into 50 µL 1X PBS to reach OD 2. Cells were spotted on poly-L-lysine coated coverslips prepared as described in [Lestini et al., 2022]. After 30 minutes incubation, coverslips were washed twice with 1X PBS. Cells were then incubated for 1 hour in Blocking solution [1X PBS – 5% Goat Serum - 0.1% Tween], then in solution containing the anti-FLAG primary antibody (Monoclonal ANTI-FLAG M2 antibody produced in mouse, Sigma, F1804) diluted 1:1000 in Antibody solution [1X PBS – 1% Goat Serum - 0.1% Tween]. Cells were then washed 5 times with Washing solution [1X PBS - 0.1% Tween], then incubated for 45 minutes in solution containing the anti-mouse secondary antibody (Horse Anti-Mouse IgG Antibody (H+L), DyLight 649, DI-2649, VectorLab) diluted 1:1000 in Antibody solution [1X PBS – 1% Goat Serum - 0.1% Tween]. Cells were then washed 5 times with Washing solution, then incubated for 10 minutes in Hoechst 33342 solution at 10 µg/mL in 1X PBS. Cells were then washed 2 times with 1X PBS, and kept in 1X PBS to be imaged.

### A.2 Microscopy

#### A.2.1 Snapshots

Snapshot imaging was performed as described in [Delpech et al., 2018]. Briefly, gfp+::rpa2+ cells cultures were inoculated overnight, then diluted to OD 0.0002, and left to grow overnight. 5 µL of culture were spotted on 1 mm thick pads of 1% agarose in 18% SW and left to dry out for 5 mins at RT. Pads were then covered with a rectangular coverslip and imaging was performed using a Zeiss Axio Observer inverted microscope with a Plan Apochromat 40x 1.4NA oil immersion objective and an AxioCam MRm camera controlled through the manufacturer’s software (ZEN 2.0). Z-stacks of 30 images with 250 nm steps and an exposure time of 150 ms were performed in the GFP channel (filter cube: excitation = BP 474/28 (HE) dichroic mirror = DFT 495 + 605 (HE), and emission = DBP 527/54 + 645/60 (HE)) and in the DIC channel for multiple position for each strain and condition.

#### A.2.2 PALM and STORM Microscopy

PALM and STORM Microscopy were performed on an IX83 Inverted microscope (Olympus) using a 100X 1.3 NA objective (Olympus) and an Orca Fusion sCMOS camera (Hamamatsu) using the Ultra-quiet read-out mode at 50 frames per second and a 2×2 binning, resulting in a pixel size of 130 nm. The microscope is controlled with micro-manager [Edelstein et al., 2014], and the microscope active z-stabilization was used to avoid z-drift. For PALM (Figure 2) we used a 532 nm laser (Voltran, 40 mW) for imaging, and a 405 nm (Voltran, 100 mW) for phototconversion a ZT532/640rpc 2-color dichroic mirror and ET600-50 (Chroma, Bellows Falls, VT, USA). 532 nm laser intensity on the sample was roughly equal to 0.1kW/cm^2^, with photoconversion intensity in the sub-W/cm^2^ range.

For STORM we used a 639 nm laser (Voltran, 140 mW), and 3 filters: a ZT532/640rpc 2-color dichroic mirror and ET700-75 emission filter, as well as an extra T660lpxr dichroic mirror (all 3 filters from Chroma, Bellows Falls, VT, USA). Laser intensity on the sample was roughly equal to 1kW/cm^2^ and 7000 images were recorded. Imaging was performed in a buffer consisting in 65 mM DABCO (D27802, Sigma), 30 mM DTT (43816, Sigma) and 30 mM Sodium Sulfite (S0505, Sigma) in TRIS-PBS pH 8.0 as in [Abdelsayed et al., 2022]. The buffer was prepared in a 300 mL batch, aliquoted in 15 mL Falcons and stored frozen at -20°C. The sample were imaged in an Attofluor imaging chamber (Invitrogen, A7816), with 1 mL of imaging buffer and another 25 mm round coverglass on top to limit air exchanges. In these conditions, FRC resolution on microtubules using similar laser intensity and the same secondary antibody was determined to be 32 nm [Abdelsayed et al., 2022]. Before STORM imaging, we record a widefield image of the DNA stained with Hoechst using a fluorescence lamp for the illumination (49000 ET – DAPI, Chroma)

Both PALM and STORM super-resolved images were reconstructed using “Detection of Molecules” (DoM2.5) [Chazeau et al., 2016], including drift-correction and grouping of consecutive localisations within a 100 nm radius. A pixel size of 20 nm was chosen for the reconstructed images used for further analysis.

#### A.2.3 spt-PALM microscopy

All spt-PALM imaging was performed using an epifluorescence microscope custom-built on a Ti Eclipse body with PFS focus stabilization system (Nikon, Düsseldorf, Germany) as previously described in [Martens et al., 2024]. The setup is equipped with five laser lines covering 405 nm, 488 nm, 638 nm, 750 nm (Oxxius, Lannion, France), and 561 nm (Novanta, Boston, USA) wavelengths which are coupled into a 70 *µ*m diameter multi-mode fibre (CeramOptec, Bonn, Germany) by a protected silver reflective collimator (RC04FC-P01, Thorlabs). Several optical elements (F950FC-A, LB1471-A-ML, LA4725-A, Thorlabs) expand the beam which is then focused with an Achromatic lens (AC254-300-A-ML, Thorlabs) and a dichroic ZT405/488/561rpc (Chroma) onto the back focal plane of a CFI Apo TIRF 60 x oil objective (NA 1.49, Nikon). To ensure uniform illumination and eliminate speckle patterns, the fiber is mechanically vibrated using a small eccentric mass motor. The heating stage was set to 45°C for all imaging experiments. Fluorescence was recorded by a Prime BSI sCMOS camera (Teledyne Photometrics, Tucson, AZ, USA; 107 nm pixel size) using ZET405/488/561m-TRF (Chroma) and bandpass filters ET510/80m (green Dendra2Hfx) and ET610/75m (orange Dendra2Hfx). The microscope setup was controlled by microManager 2.0 [Edelstein et al., 2014] combined with Pycro-Manager [Pinkard et al., 2020], while laser triggering was managed by a TriggerScope 4 (Advanced Research Consulting, Newcastle, CA, USA).

Brightfield and Pre-converted (488 nm, ET510/80m) images of the cells were first taken (10 images, 150 ms) before spt acquisitions.For spt acquisition, 15 000 images of 30 ms were taken using stroboscopic illumination of converted Dendra2Hfx (561 nm, 10 ms per frame, ET610/75m) to avoid motion blur. Photoconversion of Dendra2Hfx from green to orange was performed via primed conversion by continuous illumination with a 750 nm laser while increasing the power of a 488 nm laser strobed for 1 ms at the beginning of each frame to control conversion rate [Turkowyd et al., 2017].

### A.3 Image Processing

#### A.3.1 Widefield imaging

The widefield z-stacks were first projected onto a single image using maximum intensity projection for the GFP channel, and an average intensity projection for the DIC images. Cells were segmented from the DIC image using Cellpose (version 3.1.1.1, model cyto3 [Stringer and Pachitariu, 2025]). A threshold of 1.2x the median fluorescence in the cell is first applied, then foci are detected with a classical peak detection algorithm (relative threshold 0.7).

#### A.3.2 Spt-PALM

Localisation detection was performed with ThunderSTORM [Ovesný et al., 2014] through custom Fiji macros and red-shifted fiduciary markers (0.2*µ*m 660/60nm FluoSpheres, Invitrogen) [Balinovic et al., 2019] were used for drift correction.

Brightfield images stacks were averaged in Fiji. A custom Napari/Python (version 3.10.16) script was then used. Cell outlines were detected from brightfield images using a custom Cellpose model [Stringer et al., 2021], manually corrected, and aligned with the localisation data. Fiduciary markers localisations were automatically detected and localisations where then separated into individual cells, marker, and background. Background localisation were used to estimate noise rate for further analysis. Tracking was then performed in swift (version 0.4.3, Endesfelder et al., manuscript in prep. The swift software, as well as documentation and test data sets, can be obtained on the swift beta-testing repository upon request to the Endesfelder lab, University of Bonn.).

The achieved spatial resolution was 32.5 nm, with a field of view average between 30 and 45 nm, as obtained with Thunderstorm. A first estimate for the tracking parameters was done using Tardis [Martens et al., 2024]. Swift tracks were then used to recalculate bleach probability and expected displacement following previously published principles [Rahm et al., 2021]. Tracks with 3 or more localisations were kept for further analysis. Localisations from tracks classified as immobile in swift were then used to detect foci using HDBSCAN [McInnes and Healy, 2017, Endesfelder et al., 2013].

#### A.3.3 STORM

STORM images were analyzed using a custom Napari pipeline: first cells were segmented from the widefield Hoechst images using Cellpose (model cyto3 [Stringer and Pachitariu, 2025]), extracting general information such as cell size, shape and integrated Hoechst signal. Super-resolved foci are then segmented by applying a low intensity threshold and excluding the smallest signals, corresponding to less than 4 localisations ; the resulting foreground is transformed to a distance matrix from the background and used to find the foci centers by classical peak detection. Finally a watershed algorithm defines the region corresponding to each fluorescence foci.

The clusters of foci were identified by applying the DBSCAN algorithm [Ester et al., 1996] on detected foci centers using a neighbourhood radius (*ɛ*) of 250 nm with at least one neighbour (mim.

## Supplemental data

- Supplementary Figure 1 : Growth curves of the different strains used in this study.
- Supplementary Figure 2 : Further quantification of the widefield microscopy images.
- Supplementary Figure 3 : Quantification of the number of replication foci using widefield microscopy for GFP::RPA2 and Dendra2Hfx::RPA2
- Supplementary Figure 4 : Total number of RPA2 tracks per cell in exponential and stationary phase.
- Supplementary Figure 5 : Further quantification of the STORM microscopy images of WT cells.
- Supplementary Figure 6 : Comparison between STORM images of 45 °C and 37°C growth.
- Supplementary Figure 7 : Correlation between cell surface area and number of foci for the different conditions studied.
- Supplementary Figure 8 : Comparison between STORM images of WT and Δ*ori* cells.
- Supplementary Table 1 : Summary of the cell surface, number of cells and number of foci detected using STORM microscopy for the different strains and conditions.
- Supplementary Table 2 : Oligonucleotides used in this work.
- Supplementary Table 3 : Plasmids used in this work.
- Supplementary Table 4 : *Haloferax volcanii* strains used.

## Author contributions statement

N.O., S.D. and R.L. conceived the experiments, supervised D.N. and T.Y. D.N., R.L. and T.Y. conducted the GFP::RPA2 snapshots experiments, analyzed data, and prepared the figure. N.O and T.Y. designed and conducted PALM experiments. N.O. analyzed PALM data, and prepared the figure. T.Y. designed and conducted sptPALM experiments, developed sptPALM data analysis methods, curated and analyzed sptPALM data, and prepared visualizations. U.E. contributed to sptPALM data analysis, supervising T.Y. during data acquisition and analysis. D.N. designed and conducted STORM experiments, developed STORM analysis pipeline, and prepared visualizations. N.O., S.D. and R.L. wrote the initial draft manuscript. All authors revised the manuscript.

## Acknowledgments

The authors thank Julien Gros, Lionel Guittat, Tom Mariotte, Jean-Louis Mergny and Kate Sorg for critical reading of manuscript drafts and constructive feedback. R.L, N.O, and S.D. acknowledge recurrent funding from CNRS, Inserm, and Ecole Polytechnique as well as the Engineering for Health Interdisciplinary Center (E4H) for the New Synergies Grant Program Serge Schoen 2026. N.O. acknowledges funding from CNRS (Tremplin@INP 2021).

