## Supplemental data for "Multi-foci replication domains in *Haloferax volcanii* visualised by super-resolution microscopy"

### Supplementary data for:
### Multi-foci replication domains in *Haloferax volcanii* visualised by super-resolution microscopy

Dorian Noury, Titouan d'Yvoire, Ulrike Endesfelder, Stephane Duigou, Nicolas Olivier, & Roxane Lestini

#### Content


- Supplementary Figure 1 : Growth curves of the different strains used in this study.
- Supplementary Figure 2 : Further quantification of the widefield microscopy images.
- Supplementary Figure 3 : Quantification of the number of replication foci using widefield microscopy for GFP::RPA2 and Den-dra2Hfx::RPA2
- Supplementary Figure 4 : Total number of RPA2 tracks per cell in exponential and stationary phase.
- Supplementary Figure 5 : Further quantification of the STORM microscopy images of WT cells.
- Supplementary Figure 6 : Comparison between STORM images of 45 and 37 growth.
- Supplementary Figure 7 : Correlation between cell surface and number of foci for the different conditions studied.
- Supplementary Figure 8 : Comparison between STORM images of WT and $\Delta$*ori* cells.
- Supplementary Table 1 : Summary of the cell surface, number of cells and number of foci detected using STORM microscopy for the different strains and conditions.
- Supplementary Table 2 : Oligonucleotides used in this work
- Supplementary Table 3 : Plasmids used in this work
- Supplementary Table 4 : *Haloferax volcanii* strains used.

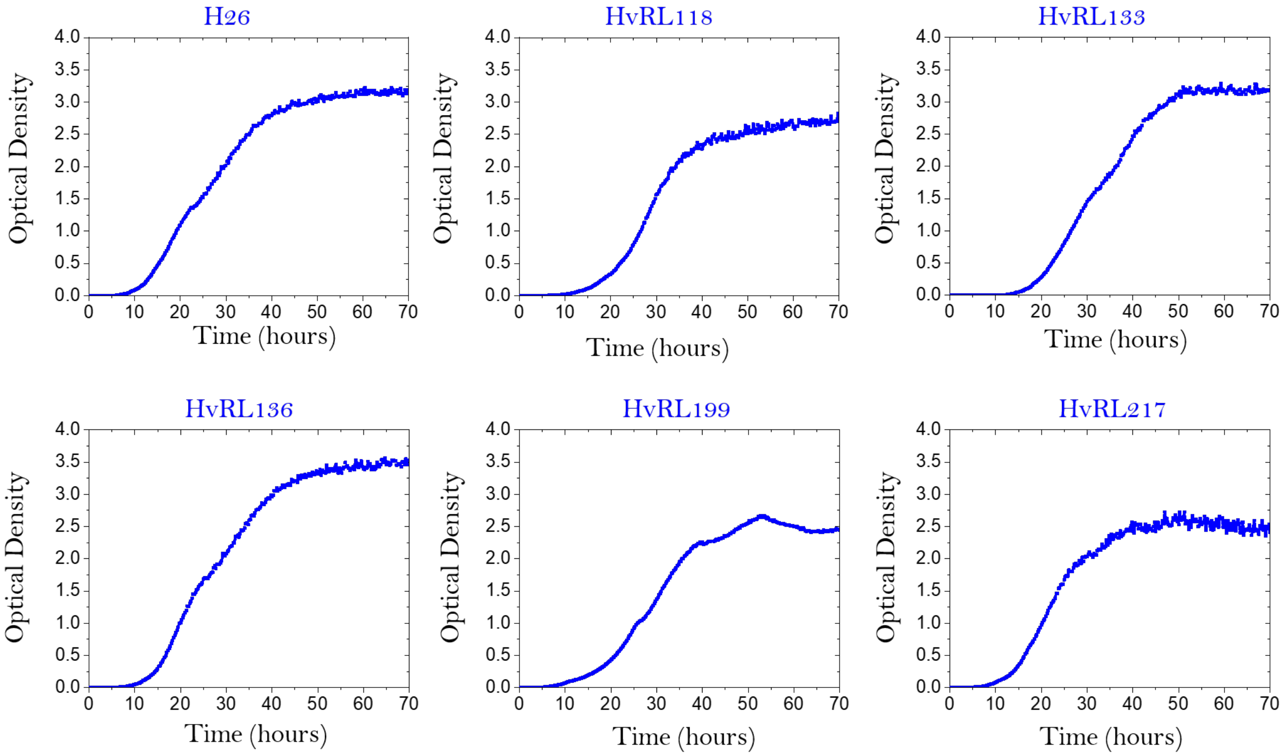


Figure S1: Representative growth curve for the different cell lines and conditions considered in the article.

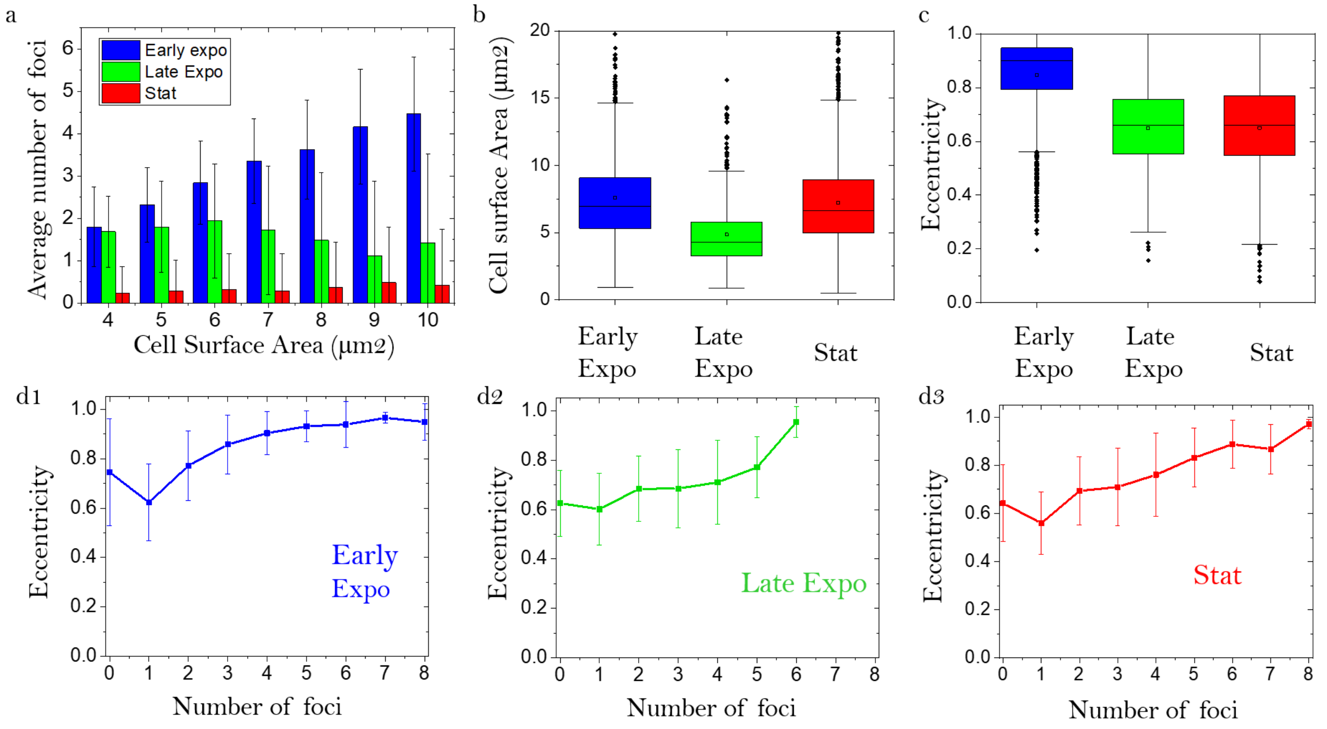


Figure S2: (a) Distribution of the average number of GFP::RPA2 foci as a function of cell surface area in different growth conditions. Bars indicate the average, and the error bars represent the standard deviation (b) Cell surface area distribution for the different growth conditions (c) Eccentricity distribution for the different growth conditions (d1-3) Average eccentricity of cells as a function of the number of detected replication foci in (d1) exponential (d2) stationary, and (d3) late exponential growth phase. (error bars represent the standard deviation)

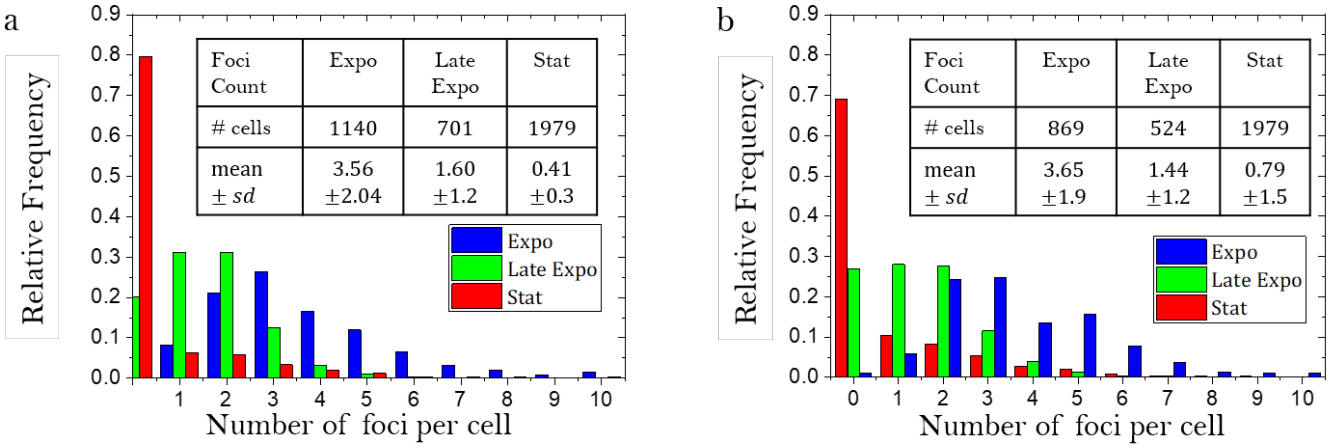


Figure S3: Comparison between the number of replication foci detected using GFP::RPA2 (a) and Dendra2Hfx::RPA2 (b) for different growth conditions

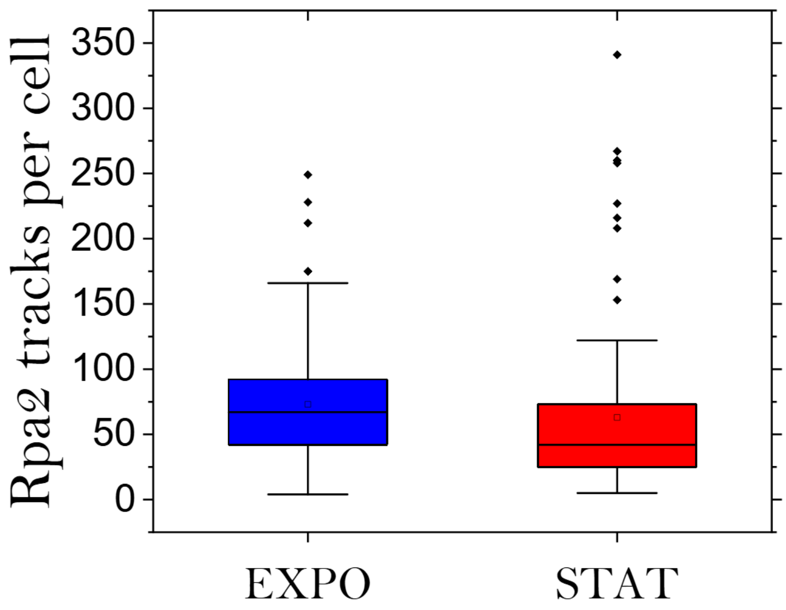


Figure S4: Total number of Rpa2 tracks per cell in exponential and stationary phase (Mann-Whitney test, p=0.0002)

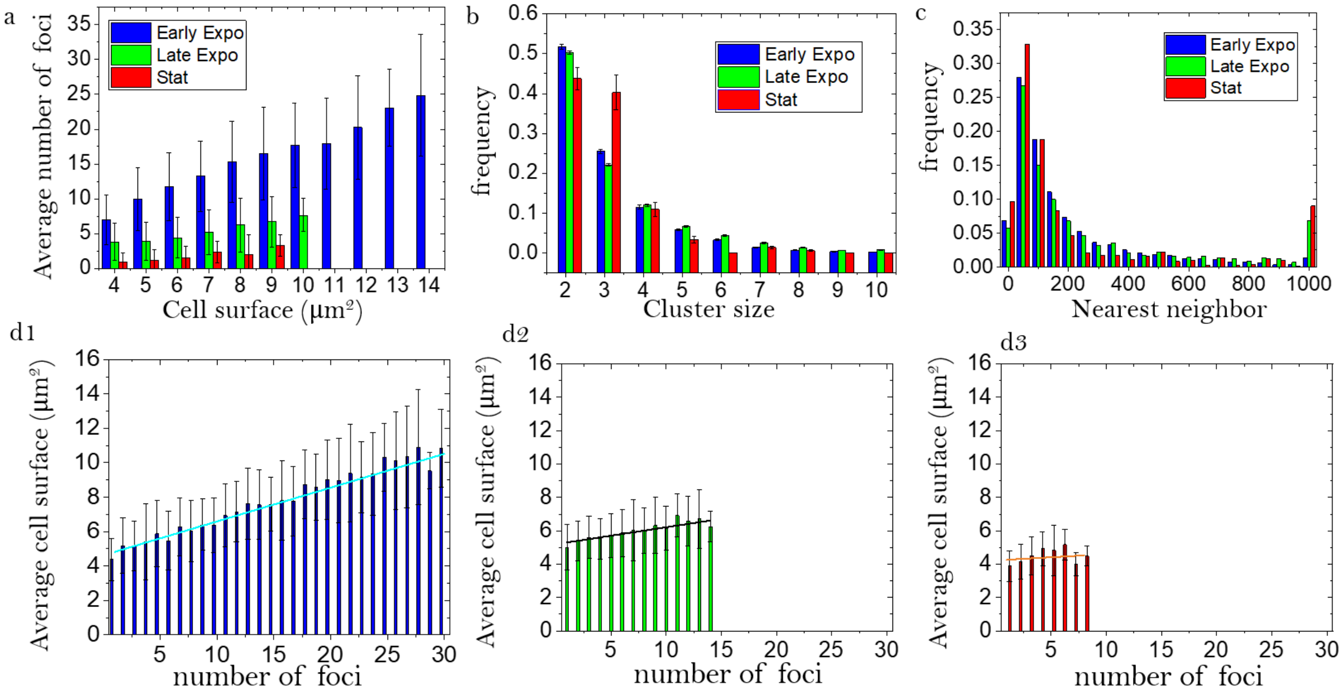


e

| Linear fit: y=ax+b | Early Expo | Late Expo | Stat |
|---|---|---|---|
| $R^2$ | 0.97 | 0.76 | -0.05 |
| Pearson coefficient | 0.98 | 0.87 | 0.23 |
| a: (Slope Surface as a function of foci) | 0.20 | 0.10 | 0.04 |
| b (µm²) | 4.6 | 5.2 | 4.2 |

Figure S5: (a) Average number of foci as a function of cell surface for early exponential, late exponential and stationary cells. (b) Number of foci per cluster (c) Distance between the nearest foci (d1-3) correlation between number of foci and cell surface for different stages of growth. (e) Summary of the data fitting from d1-d3.

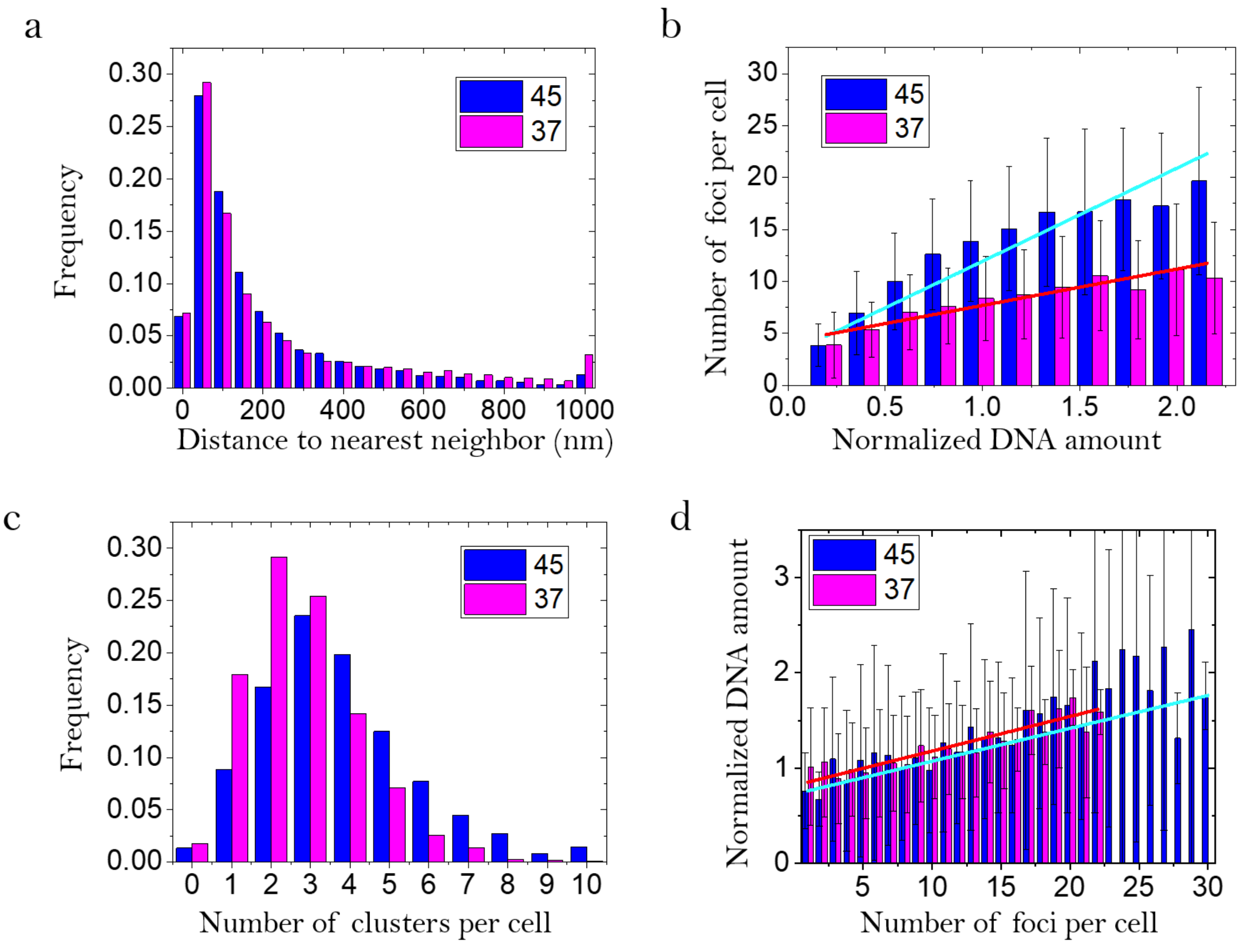


e

| | **WT 45** | | **WT37** | |
|---|---|---|---|---|
| Linear fit: y=ax+b | Foci vs normalized DNA | Normalized DNA vs foci | Foci vs normalized DNA | Normalized DNA vs foci |
| R-squared | 0.93 | 0.80 | 0.88 | 0.80 |
| Pearson | 0.96 | 0.90 | 0.94 | 0.89 |
| Slope (a) | 8.98 | 0.035 | 4.18 | 0.035 |

Figure S6: (a) Histogram of distance between the nearest foci for exponentially growing cell as 37 (pink) and 45 degrees (blue) (b) correlation between number of foci and cell surface for different temperature (c) distribution of the number of RF per cluster for both growth temperatures (d) average cell size as a function of the number of RF measured by STORM, and linear fit for both growth temperatures (e) Summary of the data fitting from (b) and (d).

a

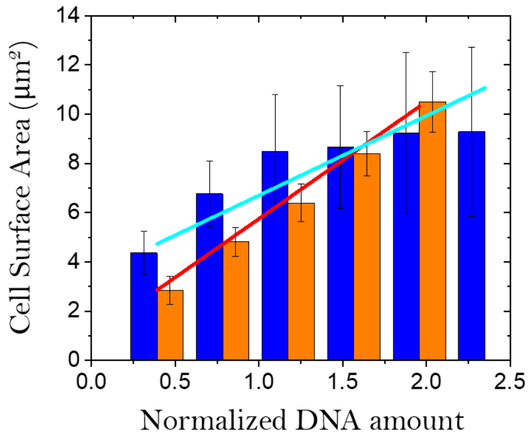


b

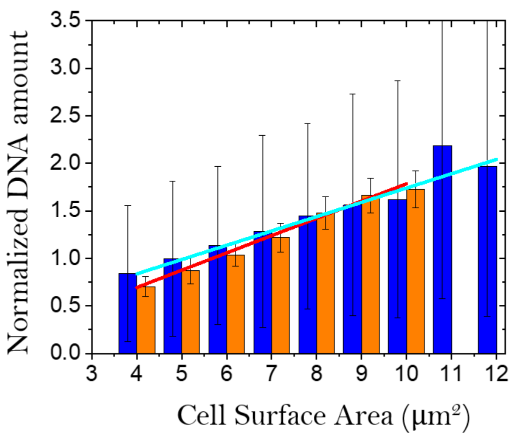


c

| | WT 45 | | Δ*ori* | |
|---|---|---|---|---|
| Linear fit: y=ax+b | Surface vs normalized DNA | Normalized DNA vs Surface | Surface vs normalized DNA | Normalized DNA vs Surface |
| R-squared | 0.86 | 0.96 | 0.99 | 0.99 |
| Pearson | 0.93 | 0.98 | 0.99 | 0.99 |
| Slope | 3.24 | 0.15 | 4.76 | 0.18 |

Figure S7: (a) Number of foci as a function of cell surface (b) cell surface as a function of number of foci and (c) table summarizing the correlation between the 2 variables

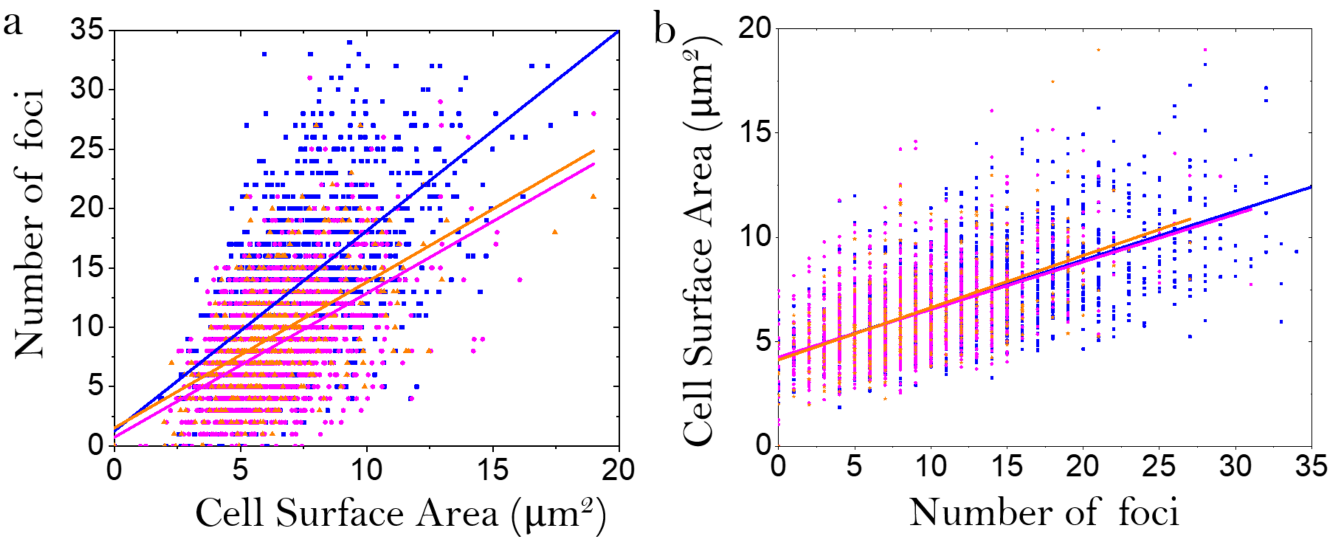


c

| | **WT 45** | **WT37** | **Δ*ori*** |
|---|---|---|---|
| Pearson Correlation coefficient | 0.63 | 0.53 | 0.55 |
| R-squared | 0.39 | 0.28 | 0.30 |
| Slope Foci as a function of surface | 1.68 | 1.21 | 1.23 |
| Slope Surface as a function of foci | 0.23 | 0.22 | 0.25 |

Figure S8: (a) Cell surface as a function of normalized DNA amount for H26 cells in blue, and $\Delta ori$ mutants in orange, with linear fit of the data in red and cyan (b) normalized DNA amount as a function of cell surface (in $\mu m^2$) for H26 cells in blue, and $\Delta ori$ mutants in orange, with linear fit of the data in red and cyan. (c) Summary of the linear fits from (a) and (b).

| Condition | Cell line | # of foci | Surface | # of cells |
|---|---|---|---|---|
| Early Expo (45°C) | WT | 13,75 ± 0,35 | 7,40 ± 0,10 µm² | 1275 (13 exp) |
| | $\Delta ori$ | 9,77 ± 0,49 | 6,56 ± 0,18 µm² | 306 (6 exp) |
| Early Expo (37°C) | WT | 8,03 ± 0,41 | 6,12 ± 0,17 µm² | 1569 (11 exp) |
| | $\Delta ori$ | 6,57 ± 0,50 | 6,17 ± 0,26 µm² | 862 (6 exp) |
| Late Expo (45°C) | WT | 4,49 ± 0,21 | 5,71 ± 0,19 µm² | 733 (4 exp) |
| | $\Delta ori$ | 3,49 ± 0,32 | 5,04 ± 0,28 µm² | 477 (3 exp) |
| Stationary (45°C) | WT | 0,84 ± 0,20 | 3,84 ± 0,25 µm² | 1233 (3 exp) |
| | $\Delta ori$ | 0,99 ± 0,11 | 4,48 ± 0,14 µm² | 689 (4 exp) |

Table S1: Summary of the cell size, surface and detected number of replication foci for wild type (HvRL118) and $\Delta ori$(HvRL136) in different growth conditions.

| Primer | Sequence (5'-3') | Relevant properties |
|---|---|---|
| pBSF2 | TTAAGTTGGGTAACGCCAGGG | Sequencing of inserts in pTA131 |
| pBSR3 | ACCCCAGGCTTTACACTTTATGC | Sequencing of inserts in pTA131 |
| RL129 | CGACGCGGAGAACTGCGCC | Screening for the presence of the *flag* sequence at the *rpa2* chromosomal locus |
| RL141 | GCCAGTGTGGACTTCGACCG | *rpa2* chromosomal locus |
| RL165 | GACGGTTACGATGTGAACTCC | *rpa2* chromosomal locus |
| RL272 | AGGTCGACGGTATCGATAAGCTTGAT ATCGCGACGCGGAGAACTGCGCCG | US with 30 bp homogy with pTA131 digested EcoRI for pRL83 and pRL118 construction |
| RL273 | GCCCTTGTCGTCATCGTCTTTGTAGT CCATCAGGCGTCACCTCCCGGAACG | US with 30 bp homogy with Rpa2' fragment containing the flag sequence for pRL83 construction |
| RL274 | CTGATGGACTACAAAGACGATGACGA CAAGGGCGTCATCCGGGAGGTCTACG | rpa2' fragment containing the flag sequence for pRL83 construction |
| RL275 | GAACAAAAGCTGGAGCTCCACCGCGG TGGCAGGTTCTCCTTCGCGCCGGCG | rpa2' fragment with 30 bp homogy with pTA131 digested NotI for pRL83 and pRL118 construction |
| RL286 | CTACAAAGACGATGACGACAAG | Screening for the presence of the *flag* sequence |
| RL287 | GTCGTCATCGTCTTTGTAGTCC | Screening for the presence of the *flag* sequence at the *rpa2* chromosomal locus |
| RL397 | CAGGCGTCACCTCCCGGAACG | US with 30 bp homogy with dendra2 fragment for pRL118 construction |
| RL398 | CGCGCGCTTCGTTCCGGGAGGTGACG CCTGATGAACACGCCGGGCATCAAC | dendra2 fragment for pRL118 construction |
| RL399 | CCAGACTTGCGACGGGAGCGG | dendra2 fragment for pRL118 construction |
| RL400 | CGATACTCGCCGCTCCCGTCGCAAGT CTGGATGGGCGTCATCCGGGAGGTC | rpa2' fragment with 30 bp homology with dendra2 fragment for pRL118 construction |
| RL401 | ACGTCAACGGCCACGCGTTCG | Sequencing |
| RL403 | ATCTCGATTCGGTGGTCGACG | Sequencing |

Table S2: Oligonucleotides used during this work.

| Plasmids | Relevant properties | Source or reference |
|---|---|---|
| pTA131 | Integrative vector based on pBluescript II, with *pyrE2* marker | Allers *et al.*, 2004 |
| pTA962-FtsZ1-Dendra2Hfx | pTA131 with *dendra2::ftsZ1* construct | Turkowyd *et al.*, 2020 |
| pRL83 | pTA131 with *flag::rpa2* construct | This study |
| pRL118 | pTA131 with *dendra2::rpa2* construct | This study |

Table S3: *Haloferax volcanii* plasmids used.

| Strain | Relevant genotype | Source or reference |
|---|---|---|
| H26 | $\Delta pyrE2$ | Allers *et al.*, 2004 |
| H1804 | $\Delta pyrE2$ $\Delta trpA$ $\Delta oriC1$ $\Delta oriC2$ $\Delta oriC3$ $\Delta oripHV4$ | Hawkins *et al.*, 2013 |
| $\Delta$ HVO_2528 | $\Delta pyrE2$ $\Delta crtI$ | Maurer *et al.*, 2018 |
| HvRL118 | $\Delta pyrE2$ *gfp::rpa2* | Delpech *et al.*, 2018 |
| HvRL133 | $\Delta pyrE2$ *flag::rpa2* | pop-in/pop-out pRL83 in H26 |
| HvRL136 | $\Delta pyrE2$ $\Delta trpA$ $\Delta oriC1$ $\Delta oriC2$ $\Delta oriC3$ $\Delta oripHV4$ *flag::rpa2* | pop-in/pop-out pRL83 in H1804 |
| HvRL217 | $\Delta pyrE2$ $\Delta crtI$ *dendra2::rpa2* | pop-in/pop-out pRL118 in $\Delta$ HVO_2528 |

Table S4: *Haloferax volcanii* strains used.
